# The geometry of pMHC-coated nanoparticles and T cell receptor clusters governs the sensitivity-specificity trade-off in T cell response

**DOI:** 10.1101/2025.02.18.638870

**Authors:** Louis Richez, Anmar Khadra

## Abstract

T cells must reliably discriminate between foreign-derived antigens that require an adaptive immune response from non-specific self-antigens that do not. This discrimination is highly specific to the affinity of the bond between ligand and T cell receptors (TCRs), as well as highly sensitive to the concentration of ligand. In this study, we examined these features of T cell mediated immunity in the context of multivalent ligand-receptor interactions between clusters of TCRs with pMHC-coated nanoparticles (NPs). Using Monte Carlo simulations of NP-T cell surface interactions, we compared the effect of TCR clustering on the dose-response profiles of various NP designs. These simulations revealed a trade-off between sensitivity and specificity, mediated by the spatial clustering of TCRs and the geometry of the NP. In particular, large clusters of TCRs were more sensitive to both NP valence and ligand concentration at the expense of antigen specificity. Conversely, uniformly distributed TCR landscapes were better suited to affinity-based ligand discrimination, while sacrificing sensitivity to ligand concentration. These features of NP-mediated T cell activation depended significantly on NP size and valence rather than on the average ligand concentration. Furthermore, we demonstrated how kinetic proofreading mechanisms may help compensate for the limitations associated with TCR clustering. These findings thus highlight the importance of interacting geometries of NP design and TCR landscape in modulating the specificity and sensitivity of the T cell response.

**Significance:** T cells rely on surface T cell receptors (TCRs) to recognize foreign antigens presented as peptide-major histocompatibility complex (pMHC) molecules. TCR clustering is crucial for T cell activation, though its full role remains not entirely clear. Using Monte Carlo simulations, we demonstrate that TCR clustering profoundly influences both surface binding dynamics of multivalent pMHC-coated nanoparticles, used in autoimmune disease therapies, as well as downstream intracellular signals leading to T cell activation. Our findings thus provide important insights into the role of interaction geometries in shaping T cell response, with implications for optimizing nanoparticle design to enhance their therapeutic efficacy.

## 1 Introduction

T cell activation is initiated by the binding of surface T cell receptors (TCRs) to specific antigen peptides associated with Major Histocompatibility Complex (pMHC) molecules. The challenge from the T cell’s perspective is to recognize specific foreign-derived antigens requiring an immune response, while tolerating non-specific self-derived antigens. T cells perform this task of ligand discrimination with remarkable sensitivity and specificity. Experiments have shown that as few as 1-10 foreign pMHC are sufficient to elicit activation of the T cell, whereas that same cell will regularly encounter thousands of similar self-pMHC without triggering a response [1–4]. Furthermore, it has also been demonstrated that the affinity of the pMHC-TCR interaction is a critical feature for the ligand discrimination task, wherein small differences in affinity can result in significant differences in the T cell response [5–7]. High sensitivity to ligand concentration and high specificity to ligand affinity are hallmarks of T cell mediated immunity [8, 9].

There is strong evidence that during T cell activation, T cell receptors and other surface proteins form micro-scale clusters that are necessary for proper signal transduction [10–14]. Other work has described that TCRs may also exist within nano-scale clusters even prior to encountering pMHC [15, 16]. It has been suggested that these TCR nanoclusters may be relevant for effective ligand discrimination by increasing the local TCR density [13], promoting cooperative binding [17] and enabling serial engagement of TCRs by single pMHC [18, 19]. Despite the TCR-pMHC interaction being itself monovalent [20], TCR clustering could result in coordinated receptor engagement and cooperative binding kinetics characteristic of polyvalent systems [21, 22].

pMHC-coated nanoparticles (NPs) have been shown to successfully stimulate T cell activation [23]. NPs coated only with disease-specific pMHCs, in the absence of any co-stimulation, have been used to promote the expansion of specific regulatory T cell populations to slow the progression of Type 1 diabetes in mice [24, 25]. These NPs benefit from being able to exploit polyvalency to enhance their avidity via multiple axes of design. For example, in addition to the affinity of a chosen ligand, the size of the NPs and the number of pMHCs per NP (also known as the valence) will also contribute to the avidity of the NPs for a given T cell surface [21, 26]. Quantifying this avidity essentially relies on knowing the number of TCRs and pMHC in the contact area as well as the monovalent pMHC-TCR affinity [21].

A large body of literature has accumulated to develop a theory describing the sensitivity and specificity of the T cell response, with particular emphasis on the intracellular signals downstream of TCR engagement. The kinetic proofreading mechanism is probably the most famous such example, and has led to many variations of the original model developed by McKeithan in 1995 [8, 27–30]. However, most of these studies investigated mechanisms of kinetic proofreading in the absence of specific geometrical considerations.

In this study, we explored how spatial receptor clustering and kinetic proofreading may work together to govern the sensitivity and specificity of the T cell response. Using a computational approach, we simulated polyvalent interactions between pMHC-coated NPs and TCR nanoclusters to examine how the spatial organization of TCRs and the design parameters of NPs interact to mediate NP avidity. Our Monte Carlo simulations captured the stochastic nature of ligand-receptor binding kinetics, the spatial organization of TCRs as well as the steric constraints of NP adsorption, revealing important trade-offs in the sensitivity of the dose-responses to various NP design parameters that depend on the precise organization of cell surface TCRs.

## 2 Results

### 2.1 TCR clusters promote polyvalent binding at the expense of NP capacity

To investigate the impact TCR clusters may have on NP avidity, we performed Monte Carlo simulations of NP binding to TCRs randomly distributed on a defined surface. These simulations took into account features of NP design, such as NP size, valence, and affinity of the ligand (Fig. 1A). We examined how these features interact with various TCR distributions on a sample cell surface of radius 1000 nm (Fig. 1B) to compare different TCR cluster sizes. We defined the surfaces by the number of TCRs belonging to each cluster. The uniform surface contained only one TCR per cluster (TPC), whereas the surface with the largest clusters contained 20 TPC (Fig. 1B). The cluster radius for each surface was defined in such a way that the intra-cluster TCR density is constant across all surfaces, with the exception of the 1 TPC (uniform) surface. While each surface was comprised of a single type of cluster, the total number of TCRs was fixed to 300 for all surfaces. Individual TCRs occupied an exclusive radius of 5 nm, accounting for the space occupied by co-receptors [12]. *[These configurations of T cell surface landscapes will be maintained throughout this study*.]

**Figure 1.**
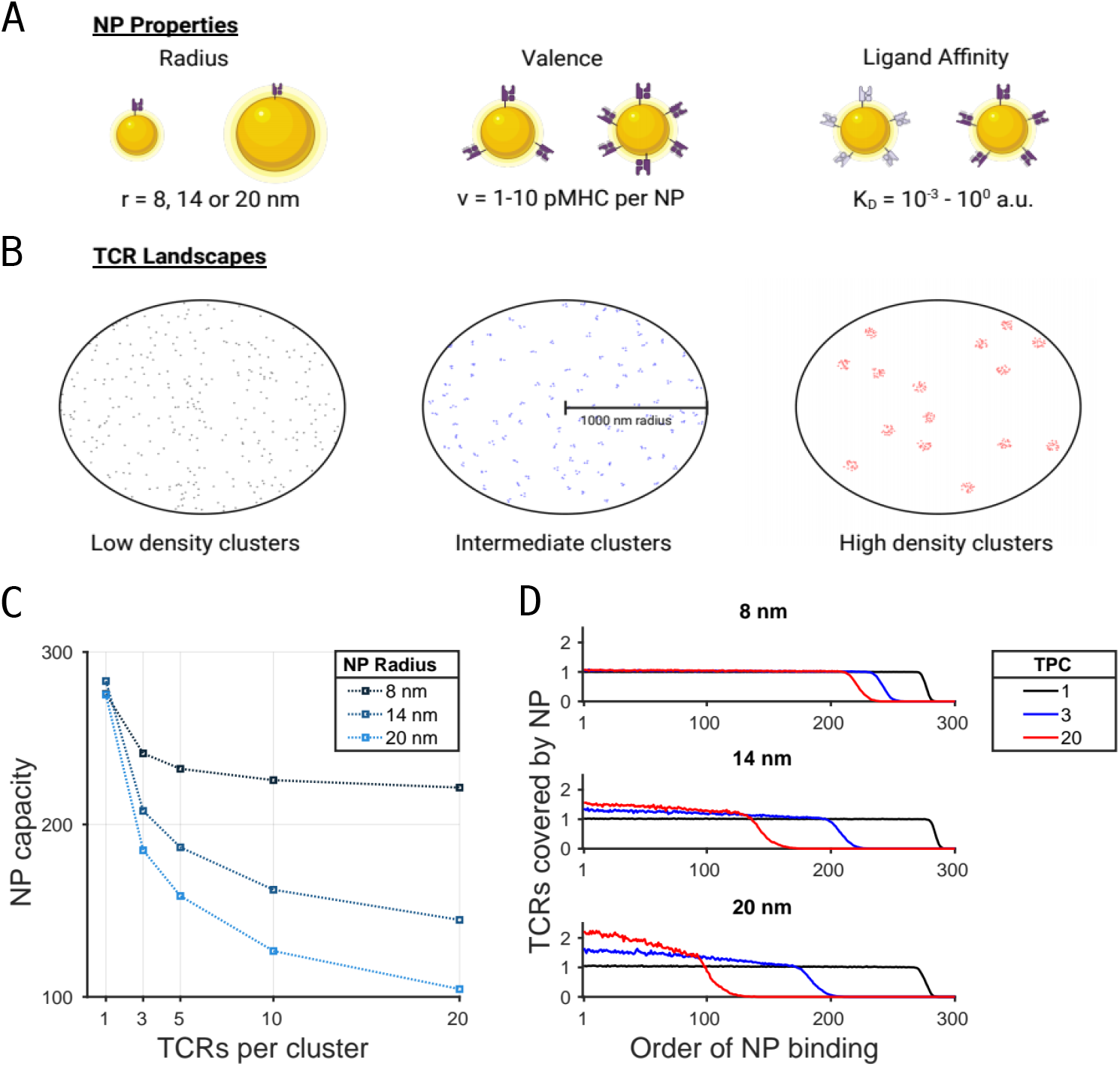
Design properties of pMHC-coated nanoparticles (NPs) and T cell receptor (TCR) landscape impose steric constraints on receptor-ligand interactions. (**A**) Simulated NPs are defined by their size, valence (i.e., number of pMHC coated on NPs), and affinity of the pMHC ligand binding to a TCR. (**B**) Three distinct TCR landscapes on a cell surface containing 300 TCRs within a 1000 nm radius. TCRs occupy a 5 nm radius and are randomly organized in clusters containing either one (left), three (center), or twenty (right) TCRs per cluster (TPC). The low density cluster (left) corresponds to uniformly distributed TCRs. (**C**) Monte Carlo estimates of NP carrying capacity of each surface in **B** for three different NP radii, averaged over 30 simulations. Curves are color-coded by NP radius (*r*) according to the legend. (**D**) Average number of covered TCRs per NP as a function of NP binding order. Each sub-panel corresponds to a different NP size with *r* = 8 nm (top), 1*r* = 4 nm (middle) or 20 nm (bottom). Curves are color-coded by type of surface landscape specified in the legend.

We defined the carrying capacity of each surface to be the greatest number of non-overlapping NPs that can simultaneously bind a given landscape. This capacity was estimated via Monte Carlo simulations by enforcing the condition that NPs remain attached to the surface provided they cover at least one TCR (essentially setting the dissociation constant to *K*_*D*_ = 0). Running these simulations until the probability of a new NP binding was less than 0.002, we then counted the number of bound NPs per surface (Fig. 1C) as well as the number of TCRs covered by each NP (Fig. 1D), averaged over 300 independent trials. Three NP sizes were considered, choosing their radii to be *r* = 8, 14 and 20 nm. As expected, larger NPs covered more TCRs at the cost of lower carrying capacity due to steric hindrance. This effect was amplified by the clustering of TCRs.

Interestingly, the carrying capacity appeared to decrease faster than the increase in covered TCRs. For example, from 1 TPC to 20 TPC, we observed a 3-fold decrease in the carrying capacity for NPs of radius 20 nm. However, this did not manifest in a 3-fold increase in covered TCRs per NP. This indicated that a significant portion of TCRs are inaccessible for binding, resulting from the inefficient packing of spheres on a plane.

While this may appear to suggest that large NPs would be a poor choice to maximize TCR engagement, this is only true once the surface is saturated with NPs. Indeed, Fig. 1D indicates that the first bound NPs cover more TCRs, facilitating polyvalent interactions with TCR clusters and thus enhancing their avidity at low NP concentrations. This motivated simulations of a wide range of NP concentrations, allowing us to generate complete dose-response profiles for the number of bound TCRs.

### 2.2 TCR clusters impact ligand discriminability in a *K*_*D*_-dependent manner

To investigate whether TCR clustering enhances ligand discriminability, we examined how variations in the dissociation constant *K*_*D*_ affect the features of the dose-response profiles. Simulations were conducted by setting the radius (*r*) and valence (*v*) of NPs to (*r, v*) = (20, 5) to ensure multivalent NP binding. These simulations were run across a range of NP concentrations for three distinct T cell surfaces: 1 TPC (uniform), 3 TPC and 20 TPC landscapes (Fig. 2A). As before, all surfaces contained the same total number of TCRs of 300, distributed in a 1000 nm radius. We considered 4 different ligand-receptor affinities defined by the dissociation constant *K*_*D*_, ranging from 10^*−*3^ to 10^0^ arbitrary units (a.u.). These values are obtained from the ratio *K*_*D*_ = *k*_off_ */k*_on_ by varying *k*_off_ and maintaining *k*_on_ = 0.1 s^*−*1^ fixed, where *k*_off_ and *k*_on_ represent the monovalent TCR-pMHC unbinding and binding rates respectively. For each condition, we performed 30 independent simulations using randomized initial TCR and NP positions. Each simulation ran for 10^6^ s of simulated time to ensure steady state distributions of bound TCRs (Fig. 2B).

**Figure 2.**
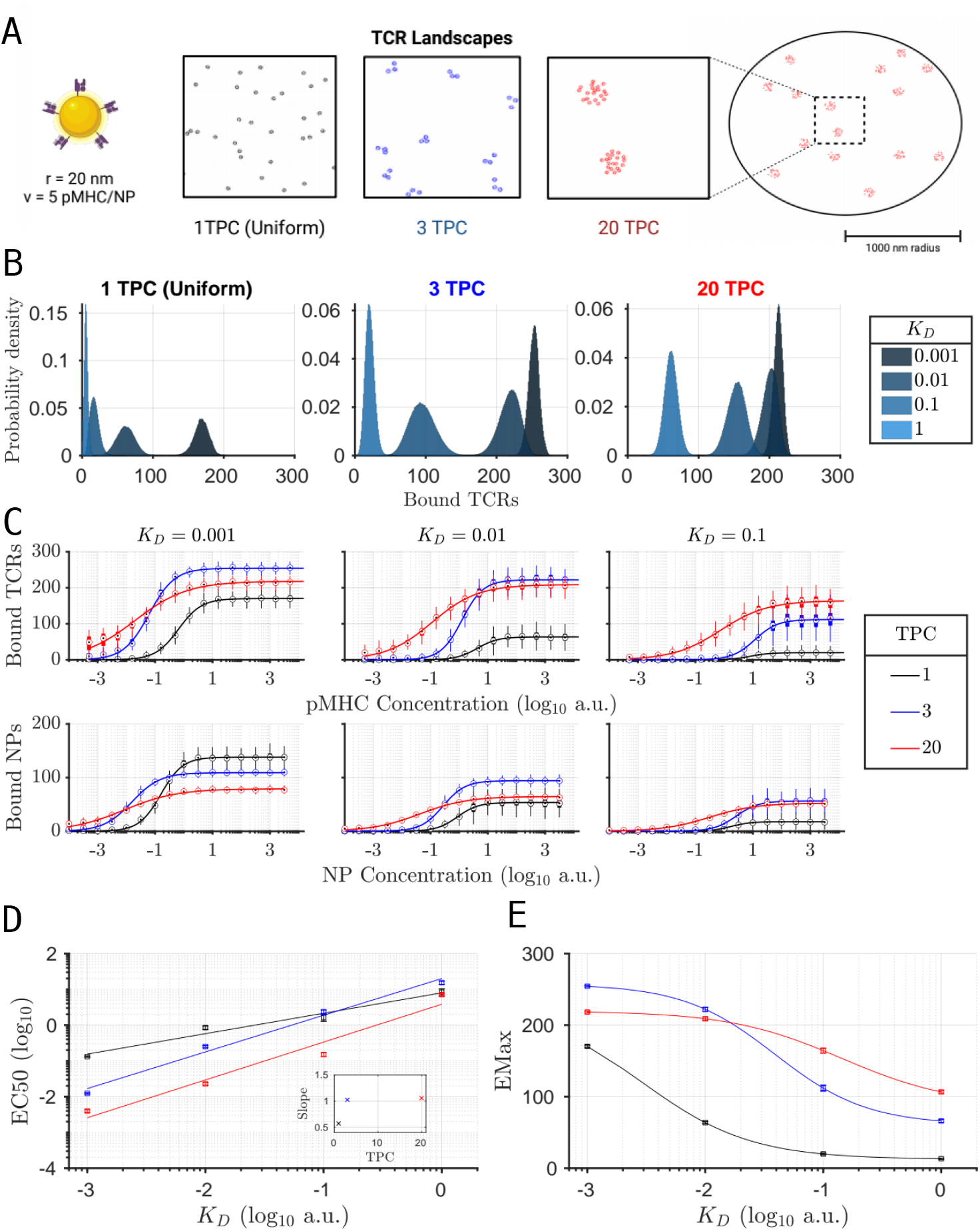
How the T cell receptor (TCR) landscape influences nanoparticle (NP) binding to surfaces. (**A**) Stochastic simulations of NPs of radius *r* = 20 and valence *v* = 5 interacting with either 1 TCR per cluster (TPC), i.e., uniform, 3 TPC or 20 TPC surfaces were performed at 16 different concentrations of NPs. Simulations were run for 10^6^ s of simulated time and repeated 30 times per condition using randomized TCR positions.(**B**) Distributions of bound TCRs of a uniform (left) and 20 TPC (right) surfaces at a fixed NP concentration (10 a.u.) for various *K*_*D*_ values.(**C**) The steady state average number of bound TCRs (top row) and bound NPs (bottom row) as a function of pMHC and NP concentrations, respectively, for three different ligand affinities: *K*_*D*_ = 0.001, 0.01, 0.1 a.u., color-coded according to the legend that specifies the number of TPC. Solid lines: Fitted Hill functions to the simulated dose-response profiles.(**D**) The EC50 of bound TCR dose-response profiles, obtained from the Hill function fits in **C**, plotted against *K*_*D*_ for each TCR landscape and color-coded according to the legend in **C**. Solid lines: The linear fits to the EC50-*K*_*D*_ plots. Inset: Slope of the linear fit for each TCR landscape. (**E**) The EMax of bound TCR dose-response profiles, obtained from the Hill function fits in **C**, as a function of *K*_*D*_ for each TCR landscape color-coded according to the legend in **C**. Solid lines: The fitted hyperbolic tangent functions to the EMax-*K*_*D*_ plots.

Our results revealed that the distributions of bound TCRs exhibit key differences between surfaces in their ability to discriminate ligands based on affinity, commonly referred to as *specificity*. This can be visually represented by the overlapping area between distributions of bound TCRs. The two highest affinity ligands, i.e. the two lowest *K*_*D*_ values, achieved the greatest separation with the 1 TPC surface (Fig. 2B). On the other hand, the 20 TPC surface was able to resolve the two lowest affinity ligands. The 3 TPC surface achieved the greatest separation between the two intermediate affinity ligands. This suggests that TCR clustering raises the threshold of *K*_*D*_ required for selective multivalent binding, promoting the avidity of low-affinity NPs.

Since these distributions only captured the response at a single NP concentration, we extended our analysis to the full dose-response profiles by considering the mean steady-state number of bound TCRs plotted against the pMHC ligand concentration (Fig. 2C). These profiles revealed that TCR clustering affects both the amplitude and the EC50 of the response, measured by the number of bound TCRs. Furthermore, the signal received by the cell did not correspond necessarily to the NP surface coverage. For example at *K*_*D*_ = 0.001, the 1 TPC surface achieved the highest level of NP binding despite generating the fewest bound TCRs. Decreasing *K*_*D*_ also caused a horizontal shift of the dose-response profiles towards lower pMHC concentrations and a vertical stretch indicating greater TCR occupancy, across all surfaces.

To further analyze the specificity of these dose-response profiles, each curve was fit to a Hill function to estimate both the EC50 and EMax. We then plotted both of these features against the dissociation constant *K*_*D*_ for all three surfaces (Fig. 2D and 2E, respectively). The average slope of the EC50-*K*_*D*_ plot, a metric commonly referred to as cooperativity, was elevated for both clustered surfaces relative to the uniform landscape; the increase occurred between 1 and 3 TPC, with little to no increase from 3 to 20 TPC (Fig. 2D, inset). Furthermore, the EC50s of the 20 TPC surface, were consistently lower than those of the 3 TPC surface. This indicates that TCR clustering enhances sensitivity by enabling detection at lower NP concentrations, facilitating an early response. The specificity of both clustered surfaces is also enhanced relative to the 1 TPC surface due to the increased cooperativity. However, larger clusters do not necessarily yield greater cooperativity, which is determined in part also by NP geometry.

It is important to note that relying on this cooperativity measure to infer ligand discriminability fails to account for the strength of the signal. Therefore, we examined how the EMax of the doseresponse curves also depend on *K*_*D*_ (Fig. 2E). For every surface, these EMax-*K*_*D*_ plots were non-linear and were all well-approximated by hyperbolic tangent functions. According to these results, we found that the presence of TCR clusters increases the amplitude of the dose-response by raising the EMax across all affinity ranges. However, larger clusters did not necessarily result in greater EMax since the 20 TPC surface had a lower capacity for NP binding due to steric hindrance compared to the 3 TPC surface. Furthermore, low affinity ligands saw the greatest relative increase in EMax as a result of TCR clustering. This indicates that TCR clusters amplify the signals from non-cognate pMHCs over those from cognate-antigens, hampering the specificity of the T cell.

We also observed that the EMax slope of the 1 TPC surface is greatest at low *K*_*D*_ and subsequently decreases for larger values, thereby suggesting that uniform TCR landscapes are sensitive to higher affinities. In contrast, the 20 TPC surface displayed greatest sensitivity at high *K*_*D*_ values. This suggests that heavily clustered surfaces are better at discriminating between low affinity ligands, whereas uniformly distributed TCRs require strong antigens to ensure sufficient TCR engagement for ligand discrimination. Small clusters containing 3 TPC, on the other hand, achieved peak specificity within an intermediate regime of ligand affinities.

These results thus demonstrate that the spatial organization of surface TCRs is critical for controlling the amplitude of the response signal. Through this control, TCR clusters can modulate both the range of affinities that yield specific binding, as well as the sensitivity to kinetic parameters. In particular, larger clusters amplify the binding of low affinity ligands but sacrifice specificity by limiting the capacity of high affinity binding.

### 2.3 Examining valence sensitivity across TCR landscapes

It has been previously demonstrated that T cell responses to NP stimulation are influenced by the valence of pMHCs displayed on NPs [23, 31–33]. In this study, we expanded this analysis by examining how NP valence impacts the specificity of binding in the context of TCR clusters. This was done by simulating the binding of *r* = 20 nm NPs to both 1 TPC (uniform) and 20 TPC surfaces (Fig. 3A) using a valence range from *v* = 1 to *v* = 10 pMHCs per NP. As before, dose-response profiles of bound TCRs were fit using Hill functions to estimate both the EC50 and the EMax under each condition.

**Figure 3.**
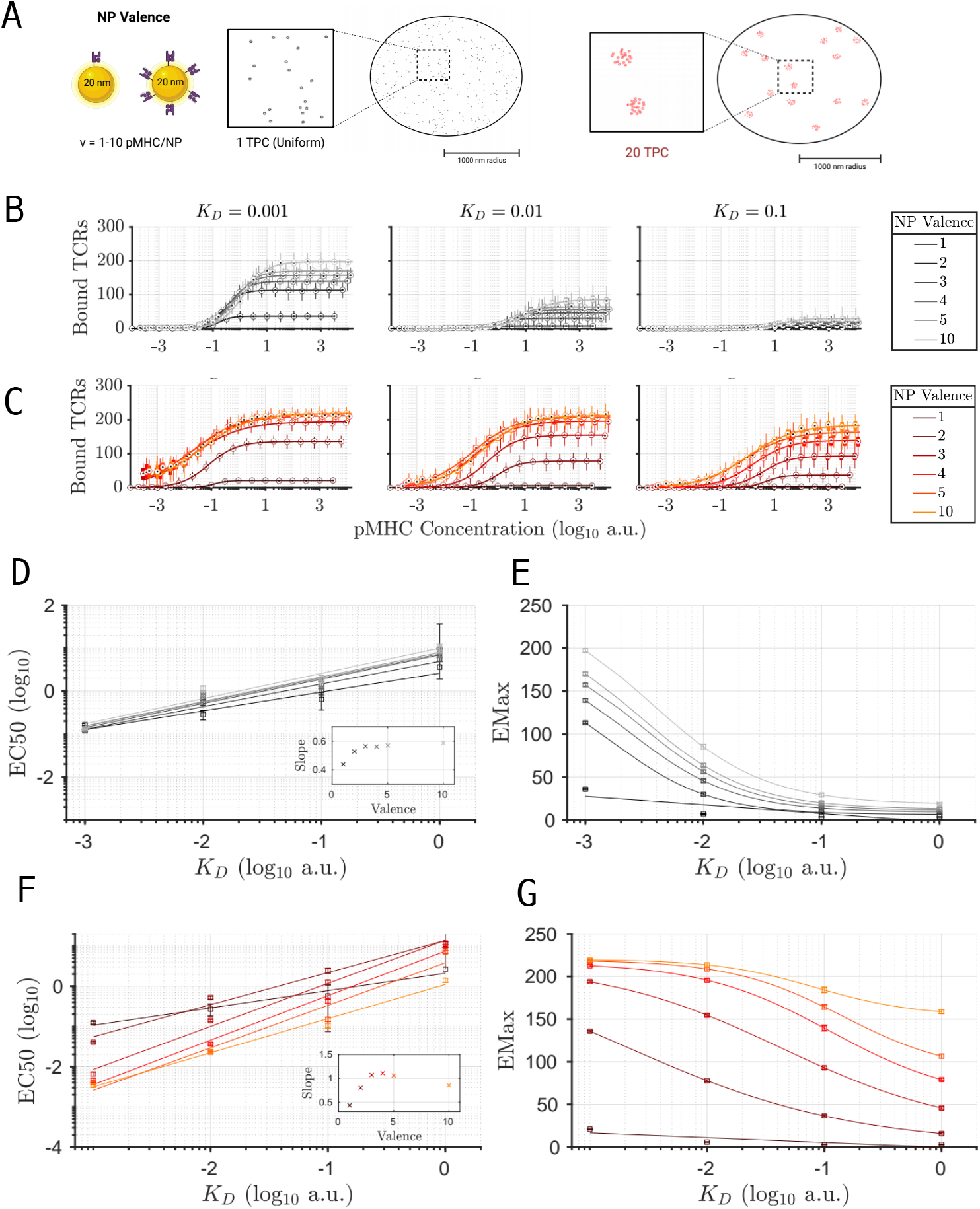
Role of valence of nanoparticles (NPs) in shaping their interaction with different T cell receptor (TCR) surface landscapes. (**A**) Schematic representations of *r* = 20 nm NPs with different valences and surfaces containing either 1 TCR per cluster (1 TPC), i.e., uniform, or 20 TPC. (**B**) Steady state average number of bound TCRs for uniform surface as a function of pMHC concentrations for three different ligand affinities (*K*_*D*_ = 0.001, 0.01, 0.1 a.u.), color-coded according to the legend that specifies NP valence (*v*). Solid lines: Fitted Hill functions to the simulated dose-response profiles. (**C**) Same as **B** but for 20 TPC surface. (**D**) The EC50 of bound TCR dose-response profiles for uniform surface, obtained from the Hill function fits in **B**, plotted against *K*_*D*_ for each NP valence and color-coded according to the legend in **B**. Solid lines: The linear fits to the EC50-*K*_*D*_ plots. Inset: Slope of the linear fit for each NP valence, a metric used to measure cooperativity.(**E**) The EMax of bound TCR dose-response profiles for uniform surface, obtained from the Hill function fits in **B**, as a function of *K*_*D*_ for each NP valence color-coded according to the legend in **B**. Solid lines: The fitted hyperbolic tangent functions to the EMax-*K*_*D*_ plots. (**F**,**G**) Same as **D** and **E**, respectively, but for 20 TPC surface, estimated from Hill function fits in **C**.

By comparing the dose-response profiles of both the 1 TPC and 20 TPC surfaces (Fig. 3B and C, respectively), we found that the 1 TPC surface is more sensitive to the affinity of the ligand rather than the valence of NPs. This was evidenced by the low level of TCR binding when *K*_*D*_ = 0.1 a.u., regardless of NP valence. However, at higher affinities, such as *K*_*D*_ = 0.001 a.u., the 1 TPC surface enabled NP discrimination based on valence. This pattern was further supported by plotting the EMax versus *K*_*D*_ (Fig. 3E), which highlighted that response amplitude is primarily determined by *K*_*D*_ in this case.

In contrast, the 20 TPC surface exhibited greater sensitivity to NP valence, particularly at low affinities (e.g., *K*_*D*_ = 0.1 a.u.). At high affinities (e.g., *K*_*D*_ = 0.001 a.u.), it became challenging to distinguish NPs with valence above *v* = 3 due to saturated levels of TCR engagement. Again, this trend was further supported by the EMax-*K*_*D*_ plots (Fig. 3G) that illustrate how response amplitude varies with *K*_*D*_. Compared to the 1 TPC condition, we observed a more pronounced separation of dose-response profiles at different valences across a wider range of affinities. At low *K*_*D*_, however, the EMax of the high valence NPs converged to the maximum TCR occupancy, causing optimal valence discrimination to occur at lower affinities.

Plotting the EC50 against *K*_*D*_ for the 1 TPC surface revealed only a modest rise in cooperativity as NP valence increased (Fig. 3D). Across all NPs, cooperativity remained below one, as measured by the slope of the linear fits, but continued to increase with the valence of the NP. The 20 TPC surface, on the other hand, conferred greater cooperativity during polyvalent interactions relative to the 1 TPC surface, as demonstrated by the steeper slopes of the EC50-*K*_*D*_ graphs (Fig. 3F). The observed cooperativity in this case was greater than one for NPs with valence between 3-5 pMHCs. Interestingly, the cooperativity appeared to peak at *v* = 4 before beginning to decline with increasing ligand number, implying that there may be an intermediate regime of NP valence that optimizes the binding specificity of the 20 TPC surface. This is also supported by the graph of EMax versus *K*_*D*_; the steepest increase in EMax was associated with a range of valences between *v* = 2 and *v* = 4.

We conclude from this analysis that surfaces with clustered TCRs are more sensitive to the valence of NPs rather than the affinity of the ligand, but the converse is true for spatially uniform TCR landscapes. Furthermore, increasing the valence of NPs always promotes ligand-discriminability in the 1 TPC case. Whereas the 20 TPC surface is most sensitive to *K*_*D*_ for an intermediate range NP valence.

### 2.4 Nanoparticle size and valence determine specificity

The size of the NPs adds another design parameter that can be leveraged to control NP avidity. On the one hand, larger NPs can cover more TCRs and can be coated with greater number of pMHCs (i.e., higher valence), thereby enhancing polyvalent binding. On the other, the surface capacity for larger NPs is reduced due to steric hindrance and competition for available space. To investigate this trade off, we simulated the interaction between NPs of radii *r* = 14 and 20 nm, coated with various levels of pMHC densities (quantified as the ratio of NP valence to NP surface area), and 20 TPC surfaces (Fig. 4A). Furthermore, we considered three separate pMHC densities: a low pMHC density of ∼ 4 × 10^*−*4^ pMHC/nm^2^ corresponding to NP parameters: (*r, v*) = (14, 1) or (20, 2); intermediate pMHC density of ∼ 10^*−*3^ pMHC/nm^2^ for NPs with (*r, v*) = (20, 5); and high pMHC density of ∼ 2 × 10^*−*3^ pMHC/nm^2^ corresponding to (*r, v*) = (14, 5) or (20, 10) NPs. We then quantified, as before, the average number of bound TCRs and NPs at steady state for each of these five NPs at three different ligand affinities: *K*_*D*_ = 0.001, 0.01, 0.1 (a.u.)(Fig. 4B).

**Figure 4.**
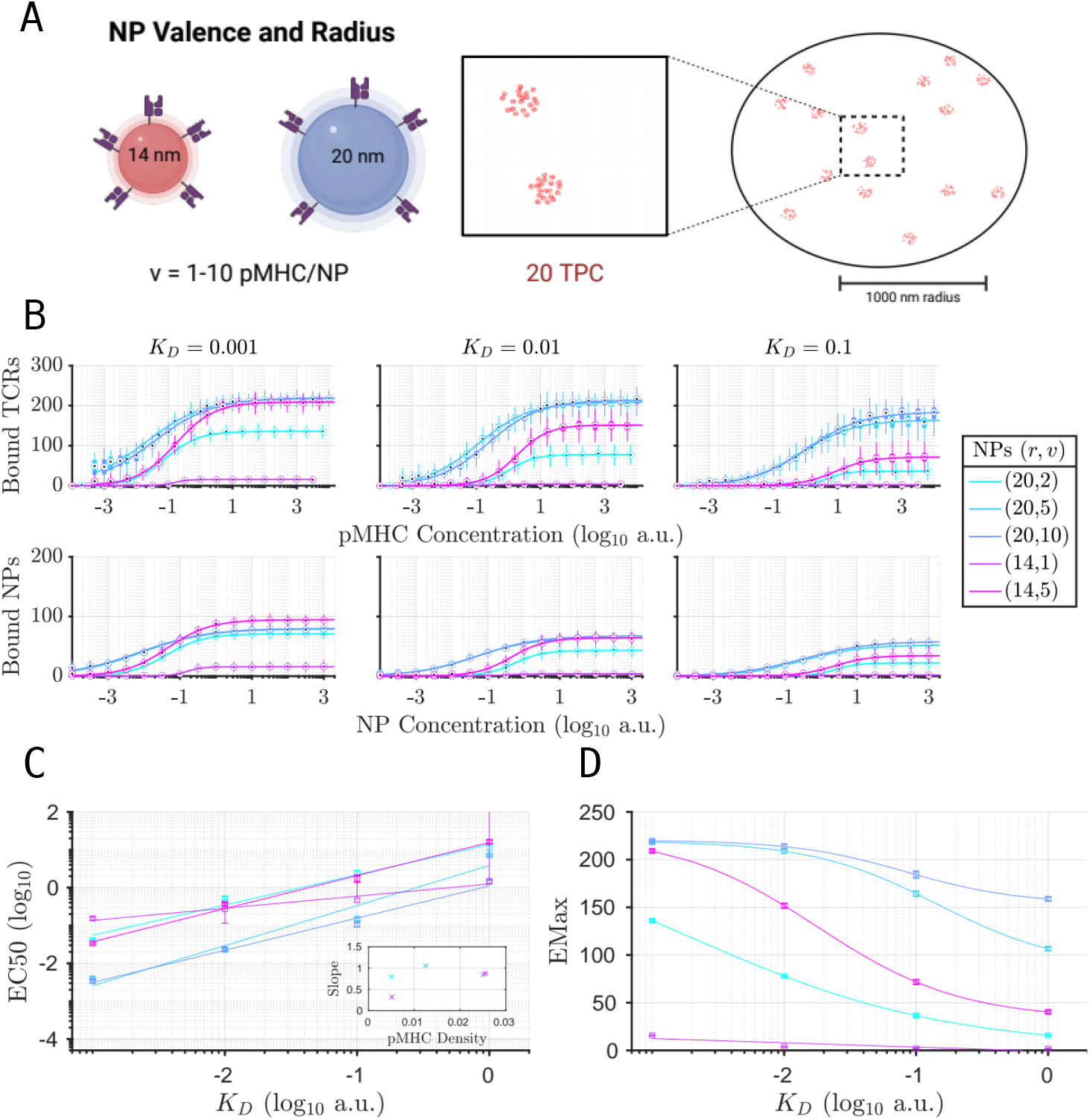
Impact of nanoparticle (NP) size on their interaction with a clustered surface containing 20 T cell receptors (TCRs) per cluster (20 TPC). (**A**) Schematic representations of NPs of different radii (*r* = 14 and 20 nm) and a 20 TPC surface. (**B**) Steady state average number of bound TCRs (top row) and bound NPs (bottom row) as a function of pMHC and NP concentrations, respectively, for three different ligand affinities (*K*_*D*_ = 0.001, 0.01, 0.1 a.u.), color-coded according to the legend that specifies the NP radius (*r*) and valence (*v*). Solid lines: Fitted Hill functions to the simulated dose-response profiles. (**C**) The EC50 of bound TCR dose-response profiles, obtained from the Hill function fits in **B**, plotted against *K*_*D*_ for each NP specified in the legend in **B**. Solid lines: The linear fits to the EC50-*K*_*D*_ plots. Inset: Slope of the linear fit for each NP specified in the legend in **B**. (**D**) The EMax of bound TCR dose-response profiles, obtained from the Hill function fits in **B**, as a function of *K*_*D*_ for each NP specified in the legend in **B**. Solid lines: The fitted hyperbolic tangent functions to the EMax-*K*_*D*_ plots.

Fitting Hill functions to the dose-response profiles of the average number of bound TCRs indicated that only the intermediate to high valence 20 nm NPs, i.e. (*r, v*) = (20, 5) and (20, 10), respectively, were able to engage more than a third of TCRs on the surface. At higher affinities, the high density (*r, v*) = (14, 5) NPs were also able to achieve comparable levels of TCR engagement, albeit with higher sensitivity to the NP concentration, as reflected by the slope of the dose-response profile.

Estimating cooperativity from the slope of the EC50 of bound TCRs versus *K*_*D*_ plot (Fig. 4C) demonstrated that (*r, v*) = (20, 5) NPs exhibited the greatest level of cooperative binding. Additionally, we found that the (*r, v*) = (14, 5) and (20, 10) NPs, which share the same pMHC density, achieved the same degree of cooperativity. However, the (*r, v*) = (20, 2) NPs displayed cooperativity twice that of (*r, v*) = (14, 1) NPs, even though they both shared approximately the same pMHC density. This indicates that cooperative binding does not uniquely depend on pMHC density, but rather necessitates polyvalent interactions within the contact area between NPs and the cell surface.

The sensitivity of EMax, derived from the Hill functions of bound TCRs, with respect to the dissociation constant *K*_*D*_ reflects how variations in ligand affinity impact TCR occupancy. Despite similar pMHC densities and similar cooperativity, the EMax of the (*r, v*) = (14, 5) NPs exhibited greater sensitivity to changes in *K*_*D*_ than the (*r, v*) = (20, 10) NPs (Fig. 4D). Both the (*r, v*) = (20, 5) and (20, 10) NPs were able to engage over a third of TCRs for all affinities tested. Their large size and high valence conferred a strong avidity, making their interactions with the 20 TPC surface relatively independent of ligand affinity, which in turn reduced the specificity of the cellular response. In contrast, the smaller (*r, v*) = (14, 5) NPs, despite having comparable pMHC density, was unable to engage as many TCRs in its contact area, especially for low affinity ligands. The EMax-*K*_*D*_ plot, in this case, showed a sharp decrease with increasing *K*_*D*_ values, indicating that the interaction of smaller NPs with the clustered surface is both highly cooperative yet highly specific.

### 2.5 Dual effects of TCR clustering and kinetic proofreading on sensitivity and specificity

TCRs undergo a number of modifications, in the form of phosphorylation steps, upon ligand binding, a well-documented process central to T cell activation [34]. The kinetic proofreading model for TCRs was developed [27] (Fig. 5A; see also Supplementary Material 5.2) and further modified to explore the impact of these phosphorylated steps on T cell sensitivity and ligand discrimination [8, 18, 27, 29, 35]. Here, we explored how the interplay between TCR clustering and kinetic proofreading affect the sensitivity and specificity of the early T cell activation signal.

**Figure 5.**
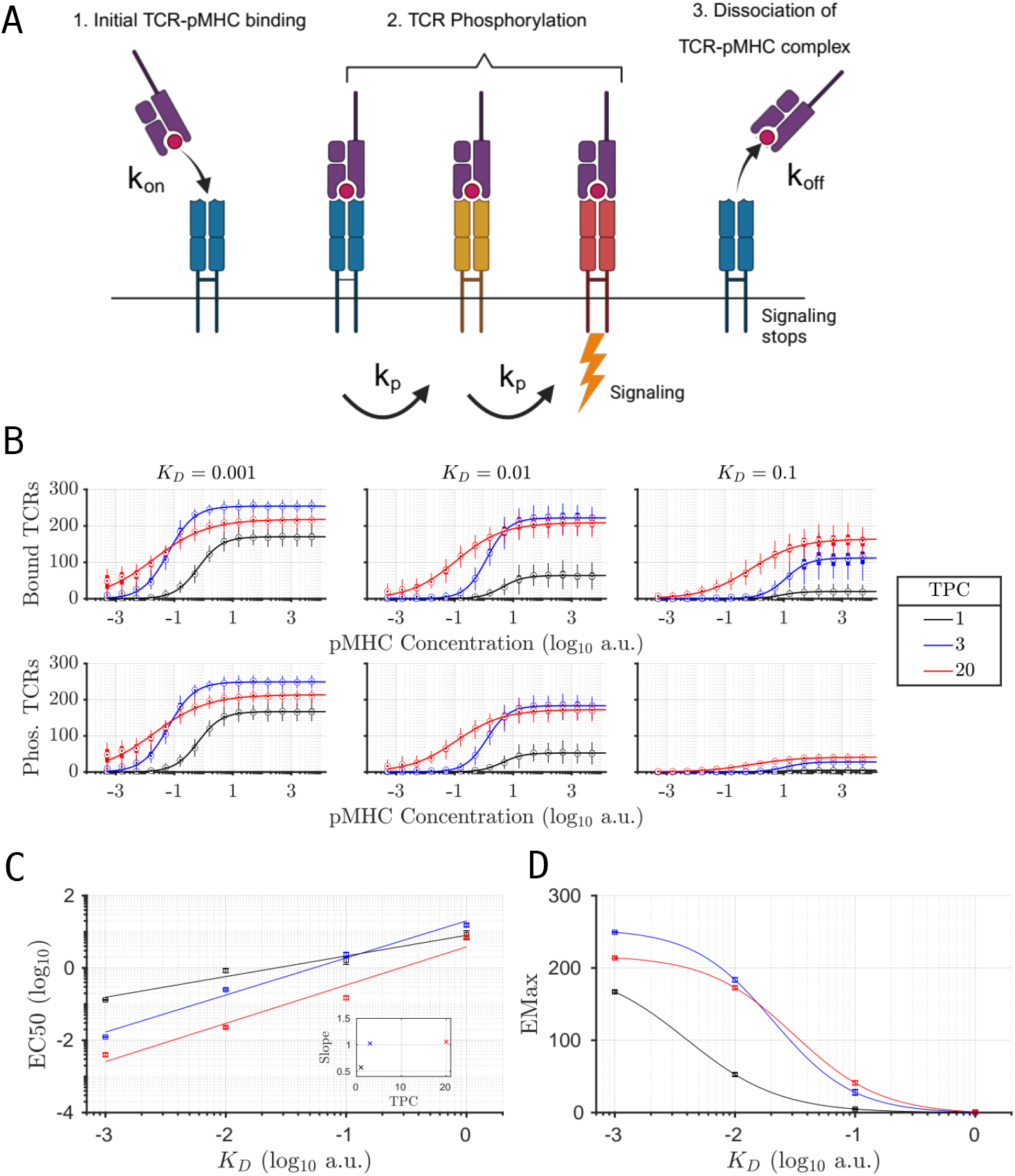
Impact of the kinetic proofreading mechanism on the sensitivity and specificity of T cells with clustered T cell receptor (TCR) surface. **(A)** Schematic of the kinetic proofreading model showcasing how the binding of a TCR to a pMHC with a rate constant *k*_on_ undergoes a series of modifications (*n* = 2) before initiating a productive signal. These modifications correspond to phosphorylation steps, each occurring with a rate constant *k*_*p*_. Dissociation of the pMHC-TCR complex with a rate constant *k*_off_ ceases signaling and reverts the TCR back to its base state. (**B**) Steady state average number of bound TCRs (top row) and phosphorylated TCRs (bottom row) as a function of pMHC concentrations for three different ligand affinities (*K*_*D*_ = 0.001, 0.01, 0.1 a.u.; *k*_on_ = 0.1), color-coded by surface type according to the legend. Solid lines: Fitted Hill functions to the simulated dose-response profiles. (**C**) The EC50 of phosphorylated TCR dose-response profiles, obtained from the Hill function fits in **B** (bottom row), plotted against *K*_*D*_ for each surface specified in the legend in **B**. Solid lines: The linear fits to the EC50-*K*_*D*_ plots. Inset: Slope of the linear fit for each surface specified in the legend in **B**. (**D**) The EMax of phosphorylated TCR dose-response profiles, obtained from the Hill function fits in **B** (bottom row), as a function of *K*_*D*_ for each surface specified in the legend in **B**. Solid lines: The fitted hyperbolic tangent functions to the EMax-*K*_*D*_ plots.

Using the canonical form of the kinetic proofreading model (Fig. 5A), it has been previously shown that the fraction of signaling versus bound TCRs [27] is given by

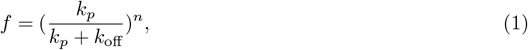

where *n* is the number of phosphorylation steps, *k*_*p*_ is the rate of each phosphorylation step and *k*_off_ is the off rate of pMHC-TCR bond. Evaluating Eq. 1 with *n* = 2, *k*_*p*_ = 0.01, *k*_on_ = 0.1 and *k*_off_ = {10^*−*4^, 10^*−*3^, 10^*−*2^, 10^*−*1^} (i.e., *K*_*D*_ = {0.001, 0.01, 0.1, 1} a.u.), we obtain the number of phosphorylated receptors by multiplying the bound TCRs from Fig. 2 by *f*. Considering the 1 TPC (uniform), 3 TPC and 20 TPC surfaces, we observed low numbers of phosphorylated TCRs for low affinity ligands despite high levels of TCR-pMHC binding for both the 20 TPC and 3 TPC surfaces (Fig. 5B). This signal attenuation was noticeably absent for the high affinity case where levels of phosphorylated TCRs were almost identical to the numbers of bound TCR-pMHC complexes, demonstrating the specificity of the kinetic proofreading mechanism even for multivalent interactions.

Interestingly, kinetic proofreading had virtually no impact on the EC50 of any dose-response curve (Fig. 5C). On the other hand, it significantly reduced the EMax amplitude of the low affinity response (Fig. 5D). This reduction was significant for both the 3 TPC and 20 TPC surface landscapes, proportional to the initial Emax of bound TCRs. As a result, the 20 TPC surface, with the highest number of bound TCRs, saw the largest decrease in amplitude at both *K*_*D*_ = 0.1 and *K*_*D*_ = 1 a.u.. In contrast, the response of the 1 TPC surface was relatively unaffected by the kinetic proofreading mechanism.

Note that, even for alternative parameter regimes of the kinetic proofreading model, we found that the 1 TPC surface was less sensitive to both variations in the phosphorylation rate *k*_*p*_ (Fig. 6A) and in the number of modifications *n* (Fig. S1 A), relative to the 20 TPC surface (Fig. 6B and Fig. S1 B). These findings suggest that the kinetic proofreading mechanism allows for the distinction of affinity from avidity in multivalent interactions, thereby enabling the T cell to benefit from the increased sensitivity granted by the clustering of TCRs without sacrificing specificity of the signal downstream of TCR binding.

**Figure 6.**
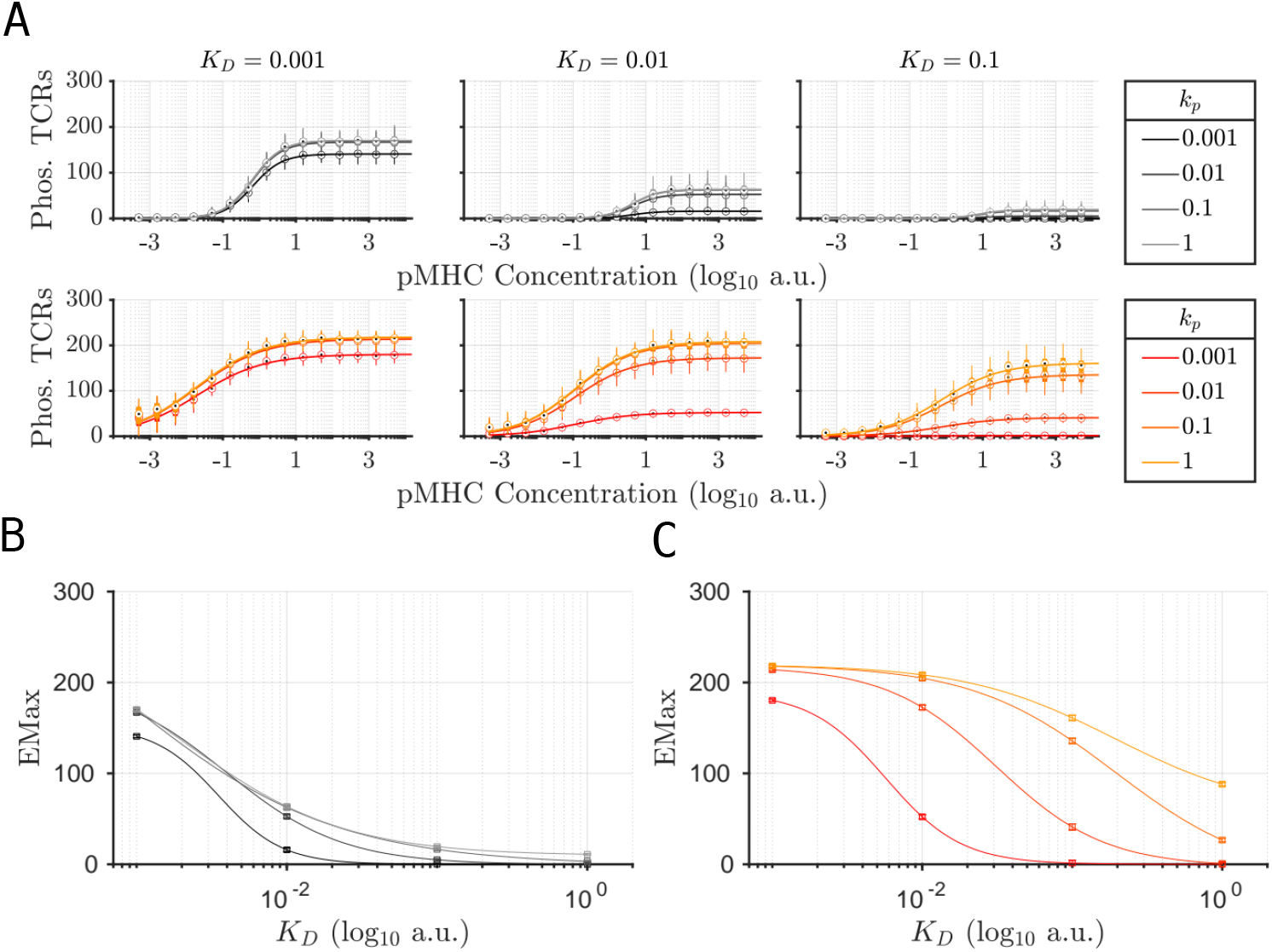
Phosphorylation rate *k*_*p*_ confers specificity by scaling EMax. **(A)** Steady state average number of phosphorylated TCRs as a function of pMHC concentrations for a 1 TCR per cluster (TPC), i.e., uniform (top row), and 20 TPC (bottom row) surfaces for three different ligand affinities (*K*_*D*_ = 0.001, 0.01, 0.1 a.u.), color-coded by the *k*_*p*_ values specified in the legend. Solid lines: Fitted Hill functions to the simulated dose-response profiles.**(B)** The EMax of phosphorylated TCR dose-response profiles of uniform surface, obtained from the Hill function fits in **A** (top row), as a function of *K*_*D*_ for each *k*_*p*_ value specified in the legend in **A** (top). **(C)** The EMax of phosphorylated TCR dose-response profiles of 20 TPC surface, obtained from the Hill function fits in **A** (bottom row), as a function of *K*_*D*_ for each *k*_*p*_ value specified in the legend in **A** (bottom).

## 3 Discussion

Our study explored the importance of TCR clustering and polyvalent interactions in mediating NP-induced T cell activation. Our model of polyvalent binding is governed by three geometric features: NP size, NP valence, and the spatial distribution of TCRs. These variables, in combination with the affinity of the pMHC-TCR complex, determine the avidity of the NP-surface interaction.

The results of our Monte Carlo simulations revealed that the interaction geometry profoundly impacts the dose-response profiles of bound TCRs as a function of pMHC concentration. In certain cases, the geometry played a more significant role than the kinetics in governing these profiles.

For example, we showed that for a fixed NP valence, the spatial organization of TCRs on the cell surface dictates the range of *K*_*D*_ values over which the cell achieves peak specificity. During activation, T cells can potentially leverage the dynamic formation of TCR clusters to resolve both ligand concentration and affinity. Furthermore, since multivalent binding is less specific to *K*_*D*_, we propose that the formation of TCR clusters during activation not only enhances greater cognate pMHC-TCR engagement, but also facilitates sampling of non-cognate pMHC concentrations. Given that elevated levels of pMHC expression are important markers of inflammation [36] and the vast majority of pMHC expressed by antigen-presenting cells (APCs) are derived from self-peptides with weak affinities [19], the clustering of TCRs following the initial engagement by high affinity cognate pMHC-TCR complexes may provide the cell with additional information about the inflammatory context. Given the vast majority of pMHC expressed on the APC surface are derived from self, the up-regulation of pMHC expression during inflammation likely results in increased concentrations of self-peptides as well. The reduced specificity due to TCR clustering can further serve to control the rise of escape mutants over the course of a viral infection, by allowing recognition of altered peptide ligands [37].

Additionally, pre-clustered TCRs on the surface may sacrifice specificity in exchange for greater sensitivity to ligand concentration and amplified responses. This trade-off would benefit antigen-experienced memory T cells by ensuring early and rapid detection at low concentrations of antigen.

On the other hand, T cell activation can benefit from the sensitivity conferred by TCR clustering while also retaining downstream specificity via an intracellular kinetic proofreading mechanism. Depending on the kinetic parameters of early TCR phosphorylation events, a small number of modifications (*n* = 2) are sufficient to filter out low affinity signals, without affecting the amplitude of the high affinity response. TCR clusters enhance the sensitivity of the T cell response by lowering the EC50 and raising the EMax of the dose-response. However, this tends to raise the EMax of the low affinity response more so than those of higher affinities. The kinetic proofreading mechanism resolves this issue by attenuating the amplitude of the low affinity signal while retaining the elevated EMax for high affinity ligands. This mechanism, combined with the cooperativity associated with clustered TCRs allows the T cell response to be highly sensitive to both ligand quality, via *K*_*D*_ specificity, as well as ligand concentration.

Our results allow us to postulate about certain design parameters of NP-based immunotherapy. Notably, the choice of NP geometry, including the valence of NPs, will determine the specificity of the treatment. Higher valence NPs will be less specific to any given TCR, thereby inducing the activation of a larger portion of the T cell population. If combined with a strong antigen, this may risk inducing a severe inflammatory response. Instead, high valence NPs can provide the necessary avidity to activate T cells using weaker antigens. This would be especially relevant in the treatment of autoimmune diseases, wherein NPs coated with self-antigens are able promote the expansion of regulatory T cells [25]. A recent study by Ols et al. has demonstrated that using multivalent antigen-coated NPs increased B cell clonotype diversity via the recruitment of low-affinity B cells, in agreement with the conclusions of our model [38]. Similar mechanisms could also be at play in the context of T cell activation, whether mediated by APCs or by NPs. Despite their low affinity, the elevated concentration of self-pMHC on the APC surface contributes to increased cell-to-cell avidity and may serve as a non-specific cue of inflammation, priming T cells for activation.

The combination of NP valence and size could also serve to target cells based on TCR organization. Since observing individual bound NPs on cell surface is easier than observing individual TCRs, this may provide an indirect method of detecting TCR clusters similar to what has been done previously [12]. This method could be expanded by using combinations of large and small NPs competing for TCRs. Leveraging the known NP properties, the number of bound NPs of each type can then be used to infer meaningful information about the size and distribution of the clusters and the density of TCRs within them.

It has been widely reported that monovalent pMHC-TCR interactions are sufficient for initiating T cell activation [3, 20]. In contrast, our simulations suggest that monovalent NPs fail to engage the number of TCRs required for robust and reliable activation. This discrepancy between our simulations and experimental observation can be partly attributed to the fact that most of these experiments rely on supported lipid bilayers (SLB) coated with either monomeric TCR or pMHC ligands. This effectively traps the receptor-ligand interaction between the cellular surface and the SLB, ensuring ample opportunities for productive TCR-pMHC encounters. In solution, T cell activation was observed with as few as 3 peptides bound to any given cell [1]; however, this result was achieved using a highly specific, strong agonist (*K*_*D*_ = 5 pM).

Finally, the conclusions of our study challenges the conventional wisdom that greater levels of polyvalent binding always results in more specific interactions [26]. Competition for TCRs, steric hindrance and the inherent stochasticity of ligand-receptor binding can place fundamental limits on the ability of T cells to resolve various features of polyvalent TCR engagement [21, 22, 39, 40]. This study highlights the critical role of TCR spatial organization in adaptive immunity and underscores the need for further experimental investigations into TCR surface arrangement. Many open questions still remain pertaining to the dynamics of TCR clustering during activation [14, 41] and the role of co-receptors and the complex spatial relationships with other surface proteins [42]. Our analysis provides a theoretical framework for the design and analysis of future such studies.

## 4 Methods

### 4.1 TCR landscapes

Surface TCR landscapes and TCR clusters were generated using the following multi-step method. First, the number of TCRs per cluster was defined; given that the total number of TCRs was taken to be 300 for all simulations, this also determined the number of clusters on the surface. The radius of the TCR clusters was set to ensure a consistent intra-cluster TCR density across surfaces. Cluster center positions were then generated using a random Poisson point process within a polar coordinate system. To ensure all clusters were fully contained within the model surface, centers were placed at least one cluster radius away from the boundary. Overlapping clusters were avoided by rejecting positions within a cluster diameter of one another, with resampling performed until the required number of non-overlapping cluster centers was achieved. Individual TCR positions within each cluster were subsequently sampled using a Poisson point process constrained by the cluster radius. To prevent TCR overlap, positions within 10 nm (this simulated TCR diameter also accounts for space occupied by co-receptors) of one another were rejected and resampled. The final TCR positions were obtained by combining the coordinates of cluster centers with those of the individual TCRs. For more details regarding the design of these TCR clusters, see Table 1

**Table 1:** Size and density of simulated TCR clusters.

| TCRs per Cluster | Number of Clusters | Size of Clusters (radius in nm) |
| --- | --- | --- |
| 20 | 15 | 50 |
| 10 | 30 | $50/\sqrt{2}$ |
| 5 | 60 | 25 |
| 3 | 100 | $50/\sqrt{6}$ |
| 1 | 300 | Undefined |

### 4.2 Stochastic simulations

Monte Carlo simulations of NP-cell surface interactions were run in MATLAB using a Tau-Leaping algorithm to model the binding kinetics [43]. This algorithm is based on the well-known Gillespie algorithm [44], but uses an adaptive time-step of length *τ*. Our implementation of this algorithm consisted of the following steps:

1. Defining the state variable *x*(*t*) = {*X*_*i*_(*t*)}, where *X*_*i*_ = −1, 0, 1, 2, 3 is the state of a single TCR (represented by *i*). Each TCR could exist as either free (unbound) with *X*_*i*_ = 0, covered (but still unbound) with *X*_*i*_ = − 1, or bound to a pMHC with *X*_*i*_ *>* 0. The bound states were further distinguished by the number of phosphorylation steps it underwent with the initial bound state represented as *X*_*i*_ = 1, the first phosphorylation step as *X*_*i*_ = 2 and the second phosphorylation step as *X*_*i*_ = 3. To reduce computational cost, only two phosphorylation steps were considered; however, adding additional steps was not expected to alter the conclusions of the kinetic proof-reading model.
2. Updating reaction rates *R*_*j*_. These rates included the propensity of TCR-pMHC binding and unbinding, phosphorylation events and NP adsorption. These reactions and their corresponding propensities are detailed in Table 2.
3. Choosing the time step *τ*. Maximum step size was set to *τ* = 500 s.
4. Generating random number of events *K*_*j*_ ∼ Poisson(*R*_*j*_*τ*) for each event type *E*_*j*_. These events correspond to the binding and unbinding of ligand-receptor complexes, TCR phosphorylation as well as the adsorption of new NPs to the cell surface. The step size was reduced if the number of reactions exceeded the current number of bound NPs. This was done to ensure efficient computations at high NP concentrations without compromising the accuracy of the steady state quantification.
5. Checking for overlap of NPs. New NP arrivals had to avoid overlap with NPs already bound to the surface. Failing this condition resulted in the removal of the new NP.
6. Updating state variables. This was done according to the equation

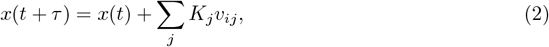

where *v*_*ij*_ represents the change on state *X*_*i*_ due to event *E*_*j*_.
7. Repeating steps 2-6, until the end of the simulation. The end was taken to be *t* = *t*_*end*_.

**Table 2:** Reactions and propensities implemented in the Tau-Leaping algorithm.

| Reaction | Propensity | Description |
| --- | --- | --- |
| NP Adsorption | $\rho K_0$ | The arrival rate of NPs on the cell surface, determined as the product of NP concentration ($\rho$) with the adsorption rate ($K_0$). |
| TCR Cross-linking | $k_{on}(v - B_{TCR})(n_t - B_{TCR})$ | The reaction that depends on the pMHC-TCR binding rate ($k_{on}$), the NP valence ($v$), the number of TCR in the contact area ($n_t$) and the current number of bound pMHC-TCR complexes ($B_{TCR}$). |
| TCR Unbinding | $k_{off}B_{TCR}$ | The product of the pMHC-TCR unbinding rate ($k_{off}$) with the number of bound TCRs ($B_{TCR}$). |
| TCR Phosphorylation | $k_p B_{TCR}$ | The product of the TCR phosphorylation rate ($k_p$) with the number of bound TCRs ($B_{TCR}$). |

Every simulation run was comprised of 30 independent trials using randomized initial conditions and TCR positions. Trials were run in parallel by means of the Parallel Computing Toolbox available for MATLAB, using servers from Compute Canada.

### 4.3 Creating and fitting the dose-response curves

The steady state number of bound TCRs was estimated from the average number of bound TCRs over the final 100 s of simulation time. Doing this across 30 independent trials provided a distribution of bound TCRs for each condition generated by perturbing a specific component of the model (e.g., TCR landscape, valence, etc.). The average of this distribution was plotted against either the NP or pMHC or concentration to produce the dose-response curves for bound TCRs or bound NPs.

These dose-response curves were then fit by Hill functions of the form

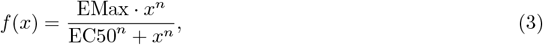

where EMax is the maximum response, EC50 is the half-maximum activation, and *n* is the Hill coefficient. The fitting was performed using the fit function from the Curve Fitting Toolbox in MATLAB to generate the best fit estimates for EMax, *n* and EC50.

Estimates for the Hill coefficient *n* would range between 0.5 and 2. We accepted non-integer estimates of *n* since this reflects the average cooperativity resulting from the stochastic ligand-receptor interactions. This is due to the random configurations of NP and TCR positions, therefore NPs will not share the exact same avidity despite being identical. Furthermore, restricting our fits to only integer values of *n* yielded poor estimates of the EC50 and EMax, which remained the primary focus of our analysis.

### 4.4 Kinetic Proofreading

The canonical kinetic proofreading (KPR) scheme consists of a number of molecular modifications to the TCR following the initial pMHC binding event (Fig. 5A). These modifications are commonly assumed to involve the phosphorylation of the TCR’s ITAMs on the cytoplasmic tail of the CD3 subunit, mediated by Zap70 and LcK [35]. These intermediate steps are critical for proper signal transduction and activation of the T cell. This mechanism has been shown to enhance the specificity of the TCR to cognate antigen at a cost to sensitivity and speed of activation.

In order to investigate how geometrical properties of NPs may affect T cell activation, we incorporated a simplified 2-step KPR scheme. Given the complexities of the signal transduction network, we make the following simplifying assumptions in our KPR model as done before [27, 35]:

- Phosphorylation steps are irreversible as long as the TCR remains bound to pMHC.
- Upon dissociation from the pMHC, the TCR instantly returns to its basal state. The TCR must undergo the complete sequence of modifications again to reach the signaling state.

### 4.5 Software and packages

Monte Carlo simulations of the NP carrying capacity as well as the analysis of the dose-response curves were performed in MATLAB R2021a using functions from the Curve Fitting Toolbox, the Parallel Computing Toolbox, the Deep Learning Toolbox and the Statistics and Machine Learning Toolbox. The stochastic simulations of NP binding dynamics for generating the dose-response curves were performed using the Compute Canada servers Narval, Cedar and Graham, running MATLAB R2018b and the afore-mentioned packages. The codes for regenerating the figures are available in Anmar Khadra’s repository: http://www.medicine.mcgill.ca/physio/khadralab/Codes/code_nanoscale2.html (uploaded Feb 18, 2025). NP and KPR schematics were made using BioRender: https://BioRender.com.

## Acknowledgments

This work was supported by the Natural Sciences and Engineering Research Council of Canada (NSERC) discovery grant (RGPIN-2019-04520) to AK. LR was supported by the NSERC-CREATE in Complex Dynamics Graduate Scholarship. The funders had no role in study design, data collection and analysis, decision to publish, or preparation of the manuscript. This research was also enabled in part by support provided by Calcul Québec (calculquebec.ca) and the Digital Research Alliance of Canada (alliancecan.ca).

## Author contributions

LR: Conceptualization, Methodology, Investigation, Writing-Original draft preparation, Visualization. AK: Supervision, Funding Acquisition, Writing-Reviewing and Editing.

## Supplementary Section

**Figure S1:**
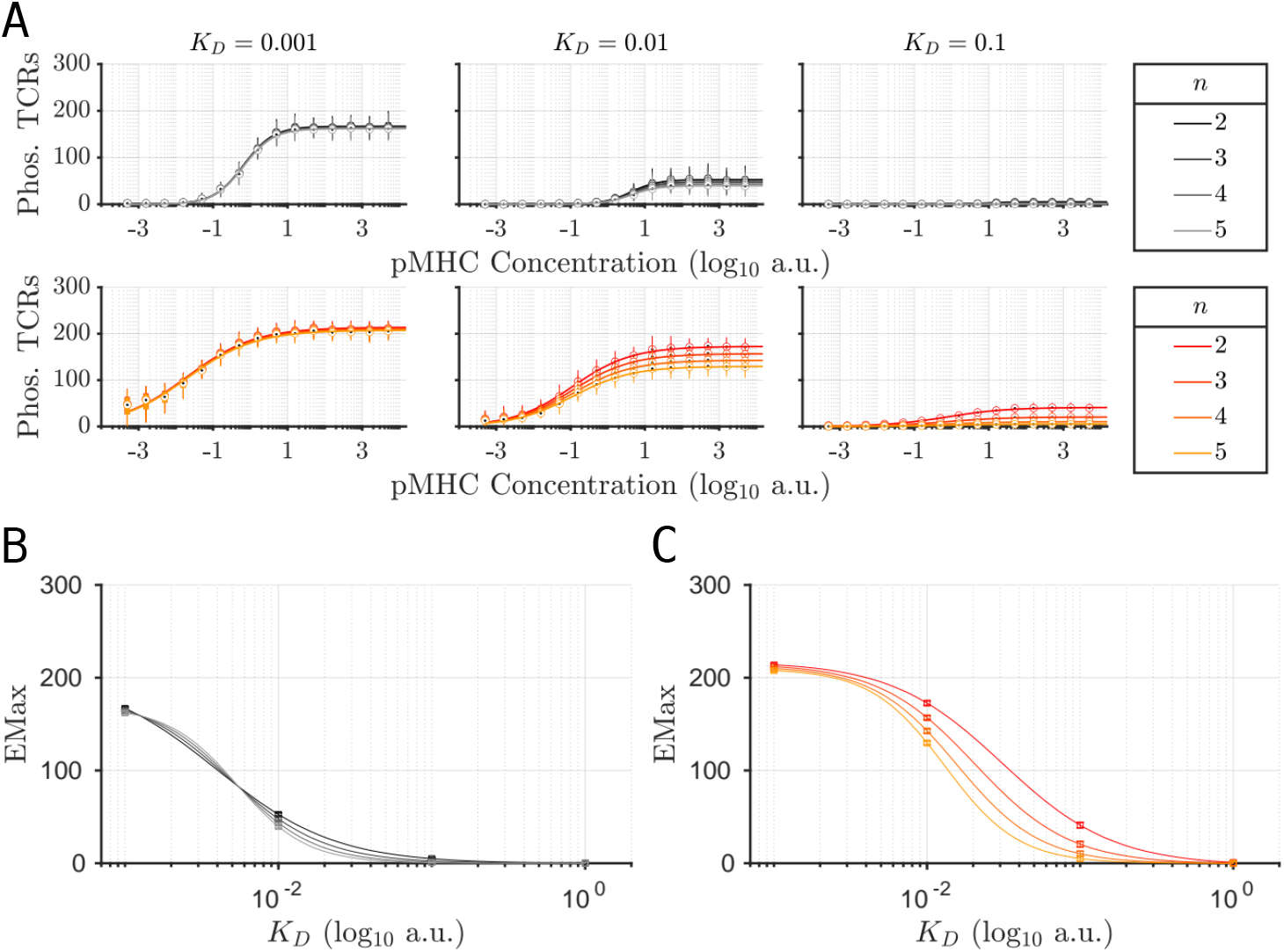
Number of phosphorylation steps *n* trades sensitivity for specificity. **(A)** Steady state average number of phosphorylated TCRs as a function of pMHC concentrations for a 1 TCR per cluster (TPC), i.e., uniform (top row), and 20 TPC (bottom row) surfaces for three different ligand affinities (*K*_*D*_ = 0.001, 0.01, 0.1 a.u.), color-coded by the *n* values specified in the legend. *k*_*p*_ = 0.01 for all curves shown here. Solid lines: Fitted Hill functions to the simulated dose-response profiles. **(B)** The EMax of phosphorylated TCR dose-response profiles of uniform surface, obtained from the Hill function fits in **A** (top row), as a function of *K*_*D*_ for each *n* value specified in the legend in **A** (top). **(C)** The EMax of phosphorylated TCR dose-response profiles of 20 TPC surface, obtained from the Hill function fits in **A** (bottom row), as a function of *K*_*D*_ for each *n* value specified in the legend in **A** (bottom).

